# Cue-reactivity and approach bias to social alcohol cues and their association with drinking in a social setting in young adults

**DOI:** 10.1101/682898

**Authors:** Martine M. Groefsema, Gabry W. Mies, Janna Cousijn, Rutger C.M.E. Engels, Guillaume Sescousse, Maartje Luijten

## Abstract

Alcohol is mainly consumed in social settings, in which people often adapt their drinking behavior to that of others, also called imitation of drinking. Yet, it remains unclear what drives this drinking in a social setting. In this study, we expected to see stronger brain and behavioral responses to *social* compared to *non-social* alcohol cues, that would be associated with actual drinking in a social setting. The sample consisted of 153 beer-drinking males, aged 18-25 years. Brain responses to social alcohol cues were measured during an alcohol cue exposure task in the scanner. Behavioral responses to social alcohol cues were measured using a stimulus-response compatibility task, providing an index of approach bias towards these cues. Drinking in a social setting was measured in a Bar-Lab setting. Specific brain responses to social alcohol cues were observed in the bilateral superior temporal sulcus and the left inferior parietal lobe. There was no approach bias towards social alcohol cues specifically, however, we did find an approach bias towards alcohol (versus soda) cues in general. Brain responses and approach bias towards social alcohol cues were unrelated and not associated with drinking, measured in the Bar-Lab. Thus, we found no support for a relation between drinking in a social setting on the one hand, and brain cue-reactivity or behavioral approach biases to social alcohol cues on the other hand. This suggests that, in contrast to our hypothesis, drinking in a social setting may not be driven by brain or behavioral responses to *social* alcohol cues.

## INTRODUCTION

Alcohol use is often initiated during adolescence and peaks in young adulthood (Chassin & Loeb, 2013; Dennis & Scott, 2007; Johnston, 2010). The health concerns associated with heavy alcohol use are manifold; including violence, sexually transmitted diseases, accidents, and increased mortality (Mokdad et al., 2018; White & Hingson, 2013). In order to prevent or reduce alcohol-related health concerns in young adults, it is important to better understand the mechanisms underlying the motivation to drink alcohol.

Drinking behavior is largely driven by social factors and alcohol is usually consumed in the company of friends, during parties and in bars (Beck et al., 2008; Clapp & Shillington, 2001; Dallas et al., 2014). Social drinking motives are often indicated as the most important reasons to drink by young adults, followed by enhancement motives (i.e., enhancing positive mood) (Kuntsche et al., 2014; Kuntsche, Knibbe, Gmel, & Engels, 2005). In line with this, it has repeatedly been found that individuals tend to adjust their alcohol consumption to a drinking partner in social settings, a phenomenon called imitation of drinking (Bot, Engels, & Knibbe, 2005; Caudill & Marlatt, 1975; Larsen, Engels, Granic, & Huizink, 2013; Larsen, Engels, Granic, & Overbeek, 2009; Larsen, Engels, Souren, Granic, & Overbeek, 2010; Larsen, Lichtwarck-Aschoff, Kuntsche, Granic, & Engels, 2013; Larsen, Overbeek, Granic, & Engels, 2010, 2012). Imitation of drinking can become problematic when individuals surround themselves with heavy drinkers as they might not be aware of it (Dallas et al., 2014). Whilst there is extensive support for imitation of drinking, individual differences in the degree to which people imitate drinking behavior are still elusive (Larsen, Engels, et al., 2013; Larsen, Engels, Wiers, Granic, & Spijkerman, 2012; Larsen, Overbeek, Vermulst, Granic, & Engels, 2010). Therefore, it is important to further examine the processes that contribute to drinking in social settings.

Previous research on heavy drinking and other drug-taking behavior has suggested that excessive drinking and the consumption of drugs is associated with the incentive value of drug-related cues, such as a glass of beer (Robinson & Berridge, 1993, 2001, 2008). Using alcohol cue-reactivity paradigms in heavy drinking and dependent samples, it was found that alcohol cues are more salient than non-alcohol cues, as they elicit responses in reward-related brain regions such as the ventral striatum (VS), ventromedial prefrontal cortex (vmPFC) and anterior cingulate cortex (ACC)(for meta-analysis see Schacht, Anton, and Myrick (2013)). Next to this process of sensitization of cues, dual process models emphasize the role of implicit cognitive biases to cues (Everitt & Robbins, 2005; Gladwin, Figner, Crone, & Wiers, 2011; Lindgren et al., 2018; Stacy & Wiers, 2010) such as approach action tendencies that can trigger alcohol use (R. W. Wiers et al., 2007; R. W. Wiers, Rinck, Kordts, Houben, & Strack, 2010). When considering drinking in a social setting, a similar process of increased incentive salience attribution and approach tendency to socially relevant stimuli might play a role in explaining individual differences in imitation of drinking. In other words, cues that have repeatedly been paired with drinking in a social setting might carry more incentive salience (i.e., motivational value) for one individual compared with another, which could eventually result in differences in the level of imitation of drinking.

To measure the salience of alcohol stimuli for young adult drinkers, who mainly drink in social settings, we used a cue-reactivity task and a behavioral approach bias task with pictures that include this social context, that is, pictures showing people having a beer/soda in a bar, in addition to plain alcohol and soda pictures. Because we expect that embedding the social setting into alcohol cues will increase their incentive value, we expect that social alcohol pictures elicit stronger brain responses than alcoholic pictures without social context in reward-related regions (e.g., VS, vmPFC, ACC), and in brain regions that are known for their role in social processing (e.g., superior temporal sulcus, temporoparietal junction, (dorso)medial prefrontal cortex, ACC) (Amodio & Frith, 2006; Apps, Rushworth, & Chang, 2016; Cousijn, Luijten, & Feldstein Ewing, 2018; Ruff & Fehr, 2014; Witteman et al., 2015). We also expect a larger behavioral approach bias to social compared to non-social alcohol cues, reflected in an approach bias to social alcohol cues. Furthermore, we expect brain cue-reactivity and behavioral approach biases to social alcohol cues to be associated with each other and with *actual* drinking behavior in a social setting. This drinking behavior was examined in the unique semi-naturalistic environment of the Bar-Lab, which has been shown to deliver an ecologically valid and informative measure of drinking behavior as the laboratory provides the opportunity to experimentally manipulate social and contextual factors without causing social desirable behavior among participants (Larsen, Engels, et al., 2013; Larsen et al., 2009; Larsen, Engels, et al., 2010; Larsen, Engels, et al., 2012; Larsen, Lichtwarck-Aschoff, et al., 2013; Larsen, Overbeek, et al., 2012). In this study we will look at 1) *imitation* of drinking (the degree to which an individual imitates the alcohol intake of his drinking partner – a confederate) and 2) *social drinking* in general (the individual’s total amount of drinks in the presence of a drinking partner).

In sum, our aim was to examine social drinking in a large group of beer-drinking young adults by triangulating three experimental measures; brain cue-reactivity to social alcohol cues, behavioral approach biases to social alcohol cues, and drinking in a social setting. We expected that heightened brain cue-reactivity and behavioral approach biases towards social alcohol cues (compared with soda cues and non-social alcohol cues) would be associated with increased drinking in a social setting. More specifically, we hypothesized that 1a) social alcohol cues would elicit more activation in reward-related and social brain regions than non-social alcohol cues; 1b) a behavioral approach bias would be stronger towards social alcohol cues than non-social alcohol cues; 2) brain cue-reactivity and behavioral approach bias towards social alcohol cues would be correlated, and 3) both measures would be positively associated with drinking in a social setting.

## MATERIALS AND METHODS

### Participants

In the context of a larger project on alcohol use in young adults (see also Groefsema et al., (2019)), participants were recruited via flyers and online advertisement. Potential participants completed an online screening to assess their eligibility to participate (see detailed flow-chart in Supplementary Figure 1). Inclusion criteria were: 1) age 18-25, 2) being male, and 3) drinking beer. Exclusion criteria were MRI contraindications and a history of brain injury. Originally, participants were further categorized into three groups-light, at-risk and dependent drinkers-based on two self-report measures collected during the initial online screening, as well as the DSM-IV (Diagnostic and Statistical Manual of Mental Disorders-5) criteria for alcohol dependence assessed later during an onsite clinical interview (Sheehan et al., 1997). The self-report measure assessed the level of alcohol-related problems (Alcohol Use Disorders Identification Test (AUDIT)) (Saunders, Aasland, Babor, de La Fuente, & Grant, 1993) and heaviness of drinking (number of alcoholic drinks per week). Yet, as we were interested in social drinking, and not heaviness of drinking, all participants were combined into one group for the current research question. For more details on our recruitment criteria, see Supplementary Figure 1. All participants participated voluntarily, gave written informed consent and received a financial compensation of 50 euros (with an additional 10 euros for the individuals who underwent an interview). The study was approved by the regional ethics committee CMO-Arnhem-Nijmegen (#2014/043).

**Figure 1:**
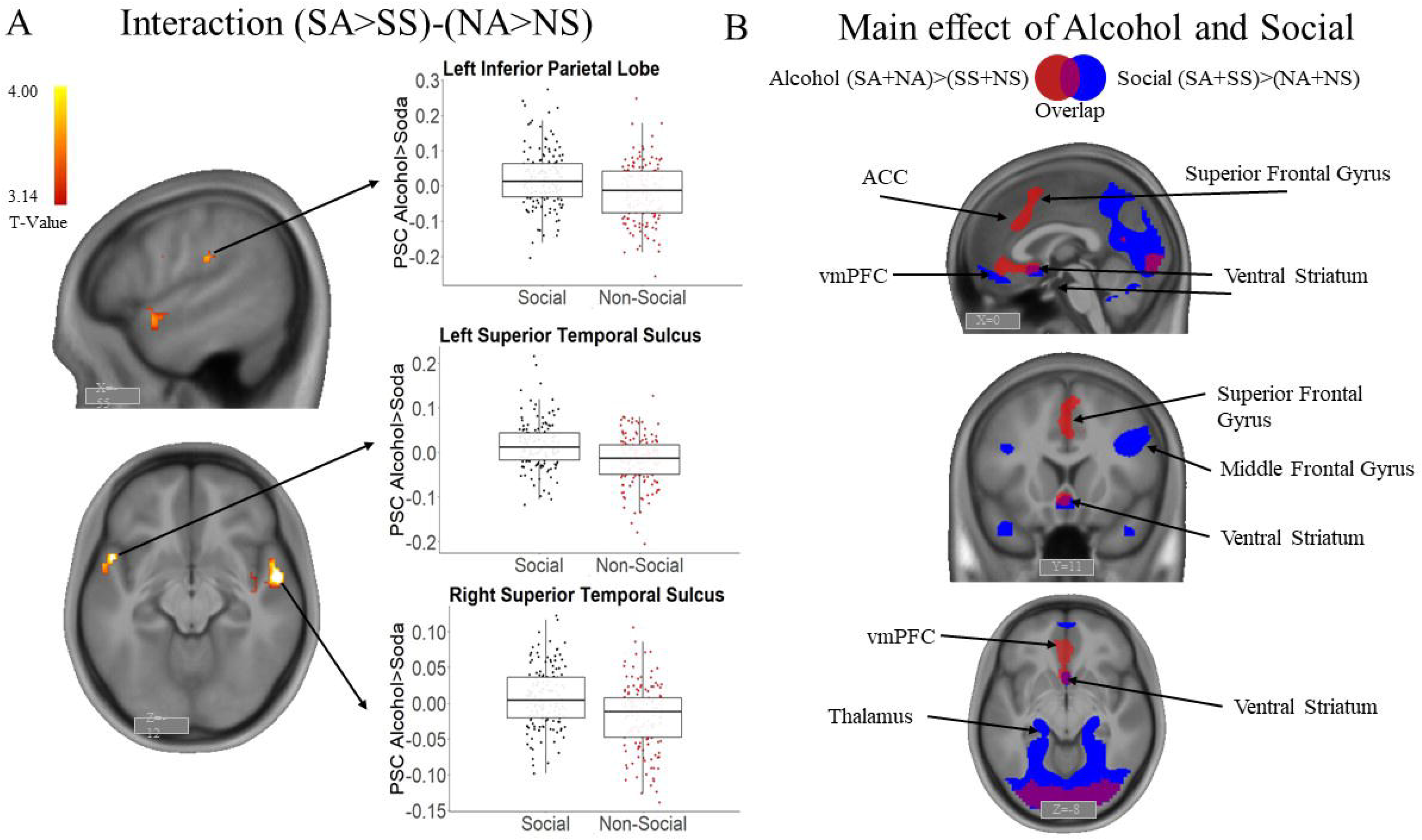
Brain responses during the Social Alcohol Cue Exposure task (A) Whole brain analysis of cue reactivity to social alcohol pictures, i.e. interaction contrast [(Social Alcohol (SA)> Social Soda (SS))-(Non-social Alcohol (NA) > Non-Social Soda (NS))]. Boxplots – reported for illustrative purposes-show the percent signal change (PSC) in the three functional clusters that show a significant interaction effect in the whole brain analysis. (B) Whole brain analysis of cue reactivity to alcohol pictures and social pictures, i.e. main effect contrasts [(Social Alcohol + Non-Social Alcohol)-(Non-Social Soda + Social Soda)] in red, and [(Social Alcohol + Social Soda) – (Non-Social Alcohol + Non-Social Soda)] in blue. Display threshold for panels A and B: voxel-level uncorrected p<.001 combined with cluster-level FWE corrected p<.05. Social= Social Alcohol-Social Soda, Non-Social= Non-Social Alcohol-Non-Social Soda, ACC = Anterior Cingulate Cortex, vmPFC = ventral medial Prefrontal Cortex.

The initial sample consisted of 166 individuals. Seven individuals were incorrectly included, as they did not meet the group criteria set out for the purpose of the broader scope of this project. In addition, six participants dropped out of the study prematurely, as they did not complete all three lab visits (see Procedure). The data of these thirteen participants were not taken into account in any of the analyses. The final sample thus consisted of 153 young adult males with a mean age of 22.78 (*SD*=1.84). They were mostly highly educated (3.3% low, 20.9% middle, 75.8% high, according to the Dutch education system), drank on average 18.09 (*SD*=13.26) alcoholic drinks per week according to the Timeline Follow-back (Sobell & Sobell, 1994), and had a mean AUDIT score of 12.69 (*SD*=6.49). Among included individuals, 17% (*n*=26) were smokers. From the final sample, slightly different numbers of participants were included in the separate analyses (see flowchart in Supplementary Figure 1; Bar-Lab measures: *n*=144, cue-reactivity task: *n*=150, and approach-avoidance task: *n*=153).

### Procedure

Following an online screening, participants completed two behavioral sessions in a Bar-Lab, followed by a separate fMRI (functional Magnetic Resonance Imaging) session, one week apart. All data collection took place between 4:00 and 10:00 pm, coinciding with typical drinking hours. Participants were asked to abstain from drinking alcohol in the 24 hours preceding testing, and sobriety was verified using a breath analyzer.

### Bar-Lab sessions

The Bar-Lab was designed to look like a real bar, increasing the ecological validity of the measure of (imitation of) drinking (Larsen et al., 2009; Larsen, Overbeek, Granic, et al., 2010). To cover the real aim of the study, participants were told that they took part in a study on the evaluation of alcohol advertisements. Participants were fully briefed after study completion. During both Bar-Lab sessions, a confederate was present, acting as a participant to facilitate imitation of drinking. Confederates were 18 males aged between 18-25, similar to the participants.

After entering the Bar-Lab, the participant and confederate were instructed to fill in online questionnaires on demographics, drinking habits, and drinking motives, followed by the rating of several non-alcoholic video advertisements in terms of attractiveness. Then, they were asked to sit at the bar, where peanuts and drinks were available, for a break lasting 30 minutes, before they had to rate video advertisements again. The experimenter offered a drink to the confederate first to set the norm and enable the examination of whether the participant would choose the same drink. Various soda drinks (200ml) and two types of local beers (250 ml, 5-5.2% alcohol) were offered. After providing the first drink, the experimenter left the Bar-Lab after explaining to the participant and confederate that they were allowed to get more drinks if they wanted to. Just before the sessions, the confederates were told to either drink one alcoholic beer followed by one soda (hereafter referred to as the “light” condition) or three alcoholic beers (hereafter referred to as the “heavy” condition) during a 30-minute ‘break’. Importantly, the confederate was instructed to initiate the drinks, by informing the participant on what he was drinking, and asking the participant if he would like something to drink as well in a neutral tone. Video and audio recordings were made during the sessions to record the number of drinks consumed. Following the break, the participant and confederate were asked again to rate the alcohol advertisements. They were also asked how they felt during the experiment, what they thought the aim of the study was (suspicion check), and how much they liked the confederate. Each participant completed a “light” and a “heavy” session with two different confederates. Session order was counterbalanced across participants and confederates. This procedure allowed us to quantify both *imitation* of drinking and *social drinking* in an ecological setting. Imitation scores were calculated by computing the difference in the number of beers consumed by the participant versus the confederate in each session, and then summing the absolute values of these differences. Social drinking scores were calculated by summing the number of beers consumed across both sessions.

### fMRI session

During the final test session, participants were instructed on the fMRI scanning procedures. The total scanning time was approximately 1 hour, during which they performed two tasks; the Social-Alcohol Cue-Exposure (SACE) task in which participants viewed alcohol-related pictures (reported in this paper) and a Beer-Incentive-Delay task (reported in (M. Groefsema et al., 2019), see Supplementary Table 1 for a complete overview of all data collected).

#### Social-Alcohol Cue-Exposure (SACE) task

We used a modified version of a passive viewing Cue-Exposure task (Schacht et al., 2011), including four conditions of interest (SA: Social Alcohol, SS: Social Soda, NA: Non-Social Alcohol, NS: Non-Social Soda, similar to Groefsema, Engels, Kuntsche, Smit, and Luijten (2016)), and one control condition (animal pictures) to which participants had to respond by a button press to ensure they paid attention to the cues. The Non-Social cues were pictures of beer or soda bottles without any human beings present, while the Social pictures showed two or more people drinking beer or soda while interacting with each other in a social setting, such as a bar. Alcohol and Soda pictures were matched one-on-one in terms of social setting, and the number and gender of people present. Twenty cues for each condition were presented in a block design: there were four epochs each consisting of four blocks with five consecutively presented pictures of the same condition (SA, SS, NA, or NS). Each picture was presented once for 4.8s. Blocks were presented in randomized order which was the same for all participants. After each block, there was a wash-out period of 6s (fixation cross) to allow for the hemodynamic response function to return to baseline. Between the four epochs, participants had a 16s break (fixation cross). Each epoch included one control cue – an animal picture presented for 4.8s to which participants had to respond – either at the beginning or at the end of a block (except for the last epoch in which 2 control cues were presented). Total task duration (including 10 practice trials) was approximately 10 minutes.

The main outcome measure of this task was brain cue-reactivity to social alcohol pictures (versus Social Soda pictures) compared with non-social alcohol pictures (versus Non-Social Soda pictures), that is, the interaction contrast ((SA>SS)-(NA>NS)).

#### fMRI data acquisition and analyses (SACE task)

Imaging was conducted on a PRISMA(Fit) 3T Siemens scanner, using a 32-channel head coil. Blood oxygen level-dependent (BOLD) sensitive functional images were acquired with a whole brain T2*-weighted sequence using multi-echo echoplanar imaging (EPI) (35 axial slices, matrix 64×64, voxel size = 3.5×3.5×3.0 mm, repetition time = 2250 ms, echo times = [9.4 18.8 28.2 37.6 ms], flip angle = 90°). The BOLD data acquisition sequence was updated during the course of the study, due to the discovery of MRI noise artifacts. The sequence parameters remained identical, except for the slice order which changed from ascending to interleaved. We took some actions in our analyses to 1) remove the artifacts, and 2) model the change in scanning sequence halfway through the study (see below). A high-resolution T1 scan was acquired in each participant (192 sagittal slices, field of view 256 mm, voxel size=1.0×1.0×1.0 mm, repetition time= 300 ms, echo time 3.03 ms).

Pre-processing steps were conducted in SPM8 (www.fil.ion.ucl.ac.uk/spm). For each volume, the four echo images were combined into a single one, weighing all echoes equally. Standard pre-processing steps were performed on the functional data: realignment to the first image of the time series, co-registration to the structural image, normalization to MNI space based on the segmentation and normalization of the structural image, and spatial smoothing with an 8 mm Gaussian kernel. In addition, two cleaning methods were incorporated into the pipeline to ensure optimal removal of artifacts and thorough de-noising of the data: 1) a Principal Component Analysis (PCA) to filter out slice-specific noise components (Viviani, Gron, & Spitzer, 2005) before pre-processing, and 2) an independent component analysis (ICA)-based automatic removal of motion artifacts using FSL (www.fmrib.ox.ac.uk/fsl) after pre-processing (ICA-AROMA; Pruim, Mennes, Buitelaar, & Beckmann, 2015; Pruim, Mennes, van Rooij, et al., 2015). This pipeline has previously been found to be efficient to take care of the MRI noise artifacts identified in the first half of our data (Nieuwhof et al., 2017).

After pre-processing, the data were modeled using a general linear model. For each condition of interest (SA, SS, NA, or NS), the various blocks of five consecutive cues were modeled as boxcars with a duration of 24s. The control condition (single animal picture) was modeled as a boxcar with a duration of 4.8s. Six motion parameters were included and a temporal high-pass filter with a cut-off of 240s (i.e., twice the maximum length between two blocks of the same condition) was applied. Parameter estimates for all conditions (i.e. SA, SS, NA, NS) were obtained by restricted maximum-likelihood estimation.

#### Stimulus-Response Compatibility (SRC) task

After scanning, approach biases were measured outside of the scanner using a well-validated Stimulus-Response Compatibility (SRC) task (Field, Caren, Fernie, & De Houwer, 2011; Groefsema et al., 2016). Participants were presented with the exact same pictures as in the SACE task (i.e., SA, SS, NA, and NS), and were instructed to either approach or avoid each picture, based on the alcohol content of the picture; ‘Approach Alcohol’ (and ‘Avoid Soda’) or ‘Avoid Alcohol’ (and ‘Approach Soda’). Every picture was presented for 2000 ms with a manikin randomly positioned above or below the picture. Participants could approach or avoid the picture by pressing the ‘up’ or ‘down’ keyboard button and thereby moving the manikin in the corresponding direction. After incorrect responses, a red cross was shown for 2000 ms, and after omissions ‘please respond faster’ was shown on the screen (2000 ms). Participants completed four blocks of 32 trials each: two blocks with only Social pictures (one with an ‘Approach Alcohol’ instruction, and one with an ‘Avoid Alcohol’ instruction), and two blocks with only Non-Social pictures (one with an ‘Approach Alcohol’ instruction, and one with an ‘Avoid Alcohol’ instruction). All pictures were presented twice, once within the ‘Approach Alcohol’ block and once within the ‘Avoid Alcohol’ block. The order of task blocks was counterbalanced across participants, with the restriction that those who started with a Social block completed both Social blocks before proceeding to the Non-Social blocks, and vice versa. Total task duration was approximately 10 minutes. The task was preceded by 16 practice trials in which participants were instructed to approach bird pictures and avoid pictures of other animals. The outcome measure of the SRC task is the approach bias in each of the four conditions (SA, NA, SS, and NS). For each condition, this approach bias was calculated by subtracting the mean reaction time observed for the “Approach” instruction from the mean reaction time observed for the “Avoid” condition including successful trials only. Errors, omissions, and outliers (responses <200 ms and >2000 ms or 3 SD above the individual mean) were discarded from these calculations (Cousijn et al., 2012; Field et al., 2011). Additionally, we calculated an interaction score by subtracting the Non-Social Alcohol bias from the Social Alcohol bias (i.e., (SA-bias>SS-bias)-(NA-bias>NS-bias)).

### Statistical analyses

All unthresholded T-maps resulting from the fMRI analyses can be accessed at https://neurovault.org/collections/IVCNOFBQ/. To test our first hypothesis (**hypothesis 1a**) regarding brain cue-reactivity to social alcohol pictures in the SACE task, a whole-brain one-sample t-test was conducted in SPM8 on the interaction contrast ((SA>SS)-(NA>NS)). Scanning sequence (i.e., pre-versus post-discovery of artifacts) was added as a binary covariate of no interest in all fMRI analyses. All T-maps were thresholded with a voxel-level uncorrected threshold of p<.001, combined with a cluster-level family-wise error (FWE) corrected threshold of p<.05, accounting for multiple comparisons across the whole brain. The probabilistic atlas of Hammers et al. (2003) was used to label significant clusters, and for visualization purposes we overlaid the output T-maps on an average T1 map of all participants, using Mango (www.ric.uthscsa.edu/mango).

To examine whether the approach bias towards social alcohol pictures was larger than the approach bias towards non-social alcohol pictures in the SRC task (**hypothesis 1b**), a 2×2 repeated-measures ANOVA was conducted, using the type of Drink (Alcohol, Soda) and Context (Social, Non-Social) as within-subject factors.

To examine the association between social alcohol cue-reactivity and social alcohol approach bias **(hypothesis 2**), a simple regression analysis was performed in SPM8 on the brain cue-reactivity interaction contrast ((SA>SS)-(NA>NS))including the interaction contrast (i.e., (SA-bias>SS-bias)-(NA-bias>NS-bias)) from the approach bias score as a covariate of interest.

To examine the relationship between the above-mentioned variables and drinking in a social setting (**hypothesis 3**), we first examined whether imitation of drinking occurred in the Bar-Lab, thereby checking the validity of this measure. For this purpose, the number of drinks in the heavy and light drinking session were compared by means of a paired-samples *t*-test. Subsequently, we included the imitation of drinking score and social drinking score as covariates of interest in two separate whole-brain analyses modelling social alcohol cue-reactivity (i.e., the ((SA>SS)-(NA>NS)) contrast). Correlational analyses were performed in SPSS (Statistical Package for Social Sciences) to examine the associations between the social alcohol approach bias (i.e., (SA-bias>SS-bias)-(NA-bias>NS-bias)) and the imitation of drinking score as well as the social drinking score.

Finally, we performed Bayesian statistics with default priors in JASP (Wagenmakers et al., 2017) in order to quantify the evidence supporting the null hypothesis. We report Bayes Factors (BF) expressing the probability of the data given H1 relative to H0 (i.e., BF_01_).

## RESULTS

### 1a Brain cue-reactivity to social alcohol pictures (SACE)

The whole-brain one-sample *t*-test on the contrast (SA>SS)-(NA>NS) revealed 3 significant clusters that survived the FWE *p*=.05 cluster-level correction: the bilateral superior temporal sulcus (STS) [x,y,z_max_= 54, −4, −11, *T*_*max*_= 4.75, k = 104 & x,y,z_max_= −51, 8, −8, *T*_*max*_= 4.48, k = 103], and the left inferior parietal lobe (IPL) [x,y,z_max_= −60, −28, 25, *T*_*max*_= 4.16, k = 66] (see Figure 1A and Supplementary Table 2). Both the main effect of alcohol cues versus soda cues (SA+NA)>(SS+NS), and the main effect of social context versus non-social context (SA+SS)>(NA+NS) revealed activation patterns in the reward-related brain network (e.g., ventral striatum and vmPFC cortex, see Figure 1B and Supplementary Table 2).

### 1b Approach bias towards social alcohol pictures (SRC)

Approach biases (i.e., faster reaction times for Approach compared with Avoid condition) occurred for all picture types, that is, all approach bias scores were significantly larger than zero (SA: *t*(152) = 12.537, *p*= <.001, SS: *t*(152) = 5.678, *p*= <.001, NA: *t*(152) = 13.612, *p*= <.001, NS: *t*(152) = 8.379, *p*= <.001). We found a stronger approach bias towards Alcohol pictures than towards Soda pictures, reflected in a main effect of Drink (*F*(1,152) = 10.639, *p*=.001, *η*^2^_*p*_ = .065 |BF_01_ = .002, with decisive evidence for alternative hypothesis). In contrast to our hypothesis, we did not find a main effect of Context (*F*(1,152) = 1.311, *p*=.254, *η*^2^_*p*_ = .009 | BF_01_=7.746, with substantial evidence for null hypothesis), nor an interaction between Drink and Context (*F*(1,152) = .335, *p*=.563, *η*^2^_*p*_ = .002 | BF_01_ = 7.111, with substantial evidence for the null hypothesis). These results suggest that the approach bias towards Alcohol pictures was similar for Social and Non-Social pictures (see Figure 2).

**Figure 2:**
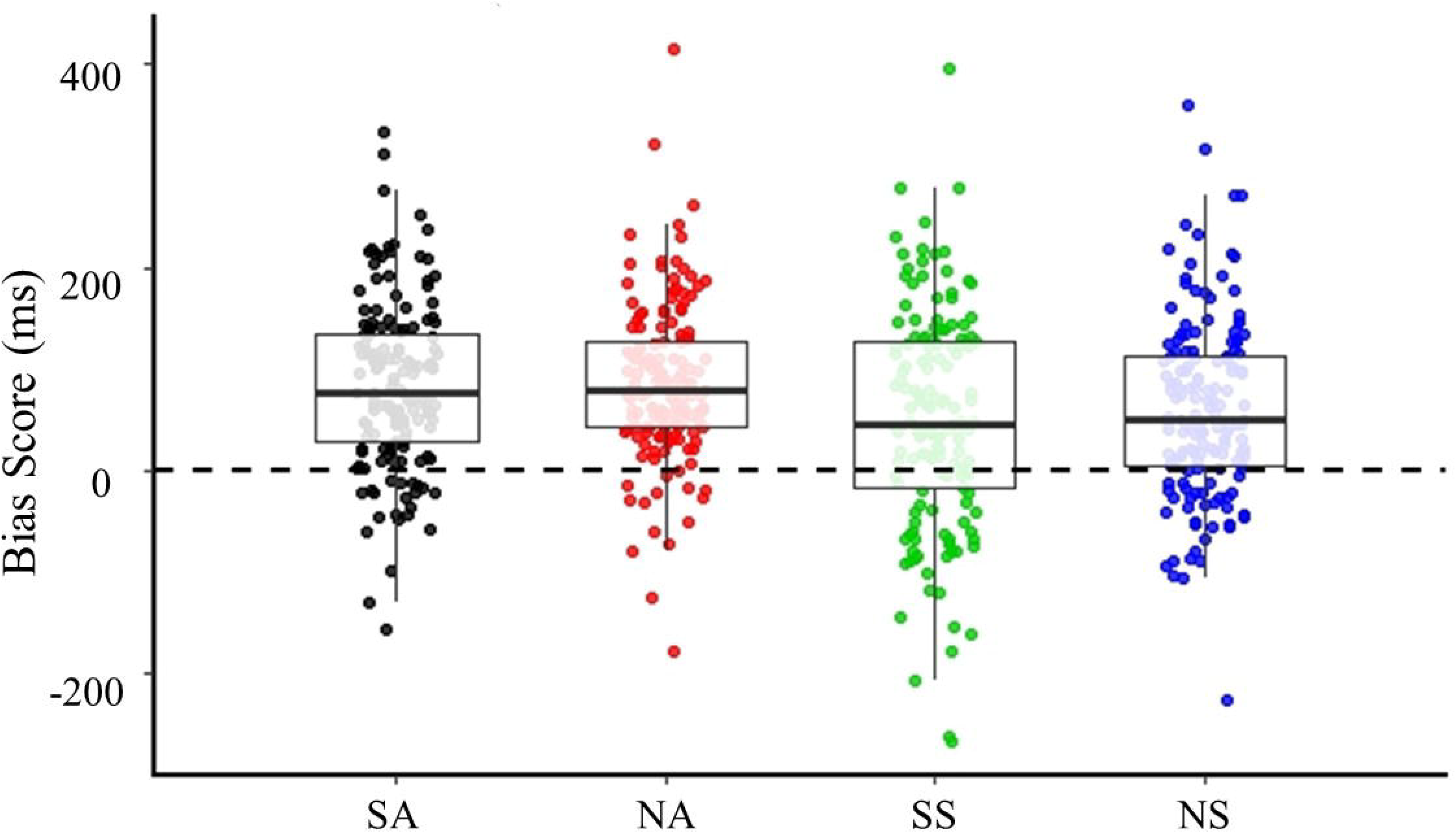
Boxplots of approach bias scores (reaction time for Avoid – Approach condition in ms) for the 4 main conditions. SA=Social Alcohol, SS= Social Soda, NA= Non-Social Alcohol, NS= Non-Social Soda. There is a significant approach bias in all conditions, as well as a main effect of Drink (p=.001), with a stronger approach bias towards Alcohol compared with Soda pictures.

### 2. Association between Social Alcohol cue-reactivity (SACE) and Social Alcohol approach bias (SRC)

The whole-brain regression analysis revealed no significant clusters for the association between social alcohol cue-reactivity and social alcohol approach bias. In addition, we performed an exploratory analysis to examine whether activation in the three regions that were originally found in the interaction effect (SA>SS)-(NA>NS) were correlated with the social alcohol approach bias. We extracted the percent signal change from these three clusters – the left and right superior temporal sulcus and the left inferior parietal lobe using MarsBaR (Brett, Anton, Valabregue, & Poline, 2002) and the rfxplot toolbox (Glascher, 2009), and regressed these values against the social alcohol approach bias scores using SPSS. We found no significant correlations (right STS r= −.154, *p*=.060 | left STS r= -.059, *p*=.471 | IPL r=−.132, *p*=.107), which was further supported by Bayesian statistics providing anecdotal to moderate evidence for the null hypothesis of no correlation between social alcohol cue-reactivity and social alcohol approach bias (right STS BF_01_ = 1.690 | left STS BF_01_ = 7.568 | left IPL BF_01_ = 2.714).

### 3. Associations between cue-reactivity or approach bias and drinking in a social setting

To check whether *imitation of drinking* occurred in our sample, we first compared the number of beers consumed by the participant in the light session with the number of beers consumed in the heavy session. As expected, participants imitated the confederate: they consumed more alcohol in the heavy session (M= 1.70, range= 0-5, *SD*= 1.26) than in the light session (M= 1.22, range= 0-4, *SD*= .96) (*F*(1,143)=19.945, *p* <.001, *η*^2^_*p*_ = .122). The imitation score, reflecting how closely participants matched the drinking pattern of the confederate, ranged from 0-4, wherein 0 reflects that the participant and confederate consumed exactly the same amount of alcohol, and higher scores reflect less imitation. The mean imitation score was 2.15 (*SD* = 1.27). The social drinking score (consumed drinks in a social drinking setting) ranged from 0-8, in which a higher score reflects more drinking during the two Bar-Lab sessions. The mean social drinking score was 2.92 (*SD* = 1.81).

Importantly, no significant correlations were found between brain reactivity to social alcohol cues and either imitation of drinking or social drinking, both at the whole brain level and within ROIs, with substantial evidence for the null hypothesis (right STS – imitation of drinking *r*=−.071, p=.391 | BF_01_ = 6.801; social drinking *r*=−.005, p=.949 | BF_01_ = 9.769; left STS – imitation of drinking *r*=−.069, p=.399 | BF_01_ = 6.879; social drinking *r*=.068, p=.409 | BF_01_ = 6.890; left IPL – imitation of drinking *r*=−.088, p=.282| BF_01_ = 5.512; social drinking *r*=−.015, p=.855 | BF_01_ = 9.627). Also no significant correlations with substantial evidence for the null hypothesis were found between the social alcohol approach bias score and either imitation of drinking, (*r*=.035, *p*=.680 | BF_01_ = 8.816) or social drinking (*r*=−.005, *p*=.956 | BF_01_ = 9.577).

Additionally, we performed exploratory analyses to examine whether alcohol cue-reactivity and alcohol approach biases, independent of social context ((SA+NA)-(SS+NS)), showed correlations with each other and with imitation of drinking or social drinking. Using a whole-brain simple regression model, we found no significant correlation between alcohol cue-reactivity and alcohol approach bias. Furthermore, we did not find any significant correlations between alcohol cue-reactivity and imitation of drinking or social drinking at the whole-brain level, or between the alcohol approach bias scores and imitation of drinking or social drinking.

## DISCUSSION

This study examined the relationship between brain and behavioral responses to *social* alcohol cues, as well as how these measures relate to drinking in a social setting. We included a large sample of young adults and measured imitation of drinking using a semi-naturalistic Bar-Lab setting. First, we observed brain reactivity specifically towards *social* alcohol cues in the bilateral superior temporal sulcus and the left inferior parietal lobe, as indicated by stronger responses to social versus non-social alcohol cues, compared with social versus non-social soda cues. Second, we found no support for an approach bias towards *social* alcohol cues specifically, however, we did find an approach bias towards alcohol (versus soda) cues independently of social context. Third, brain reactivity and behavioral approach bias towards *social* alcohol cues were uncorrelated with each other, and were neither related to imitation of alcohol use or social drinking (for an overview of all findings see Figure 3). Additional exploratory analyses showed that, regardless of social context, brain and behavioral responses to alcohol cues were similarly uncorrelated with each other, and were also not correlated with drinking in a social setting.

**Figure 3:**
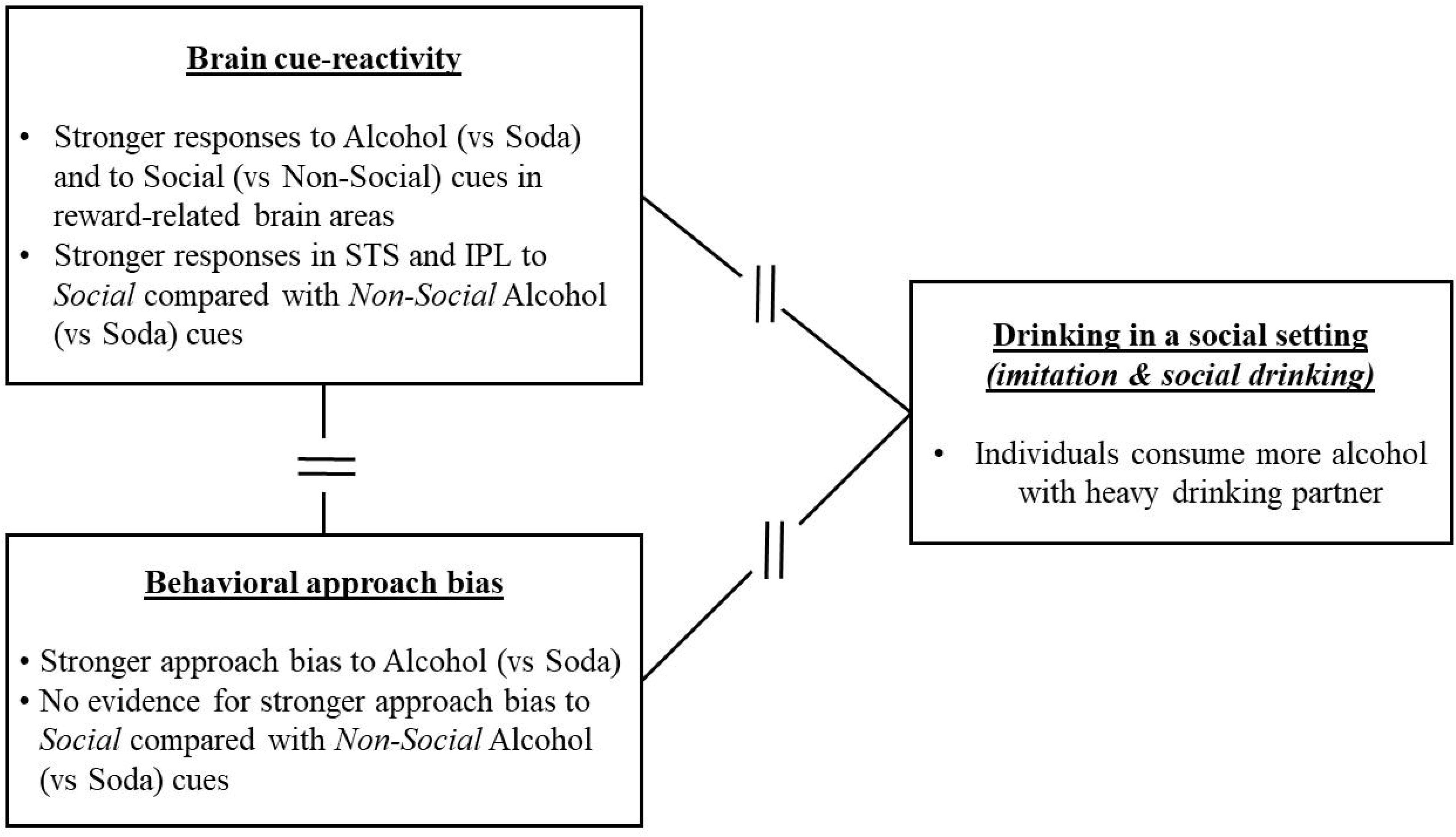
Overview of the results. Broken lines reflect no significant associations between the variables. STS = Superior Temporal Sulcus, IPL = Inferior Parietal Lobe

Since the social drinking setting is thought to be highly salient for young adult drinkers, we were interested in the effect of social context on alcohol cue-reactivity. At the brain level, against our expectations, we found no increased activation in reward-related brain regions, such as the ventral striatum, in response to social versus non-social alcohol cues. Instead, increased activation was found in the superior temporal sulcus (STS) and the left inferior parietal lobe (IPL). Whilst these brain areas have not often been emphasized in individual cue-reactivity studies, two meta-analyses performing activation likelihood estimation (ALE) analyses across these studies revealed an association between parietal lobe activation and craving (Chase, Eickhoff, Laird, & Hogarth, 2011), as well as stronger brain responses in the superior temporal sulcus to alcohol versus neutral cues in alcohol-dependent patients versus healthy controls (Schacht et al., 2013). Interestingly, the temporal pole has also been implicated in emotion processing (Olson, Plotzker, & Ezzyat, 2007) and social cognition (Adolphs, 2009; Allison, Puce, & McCarthy, 2000; Insel & Fernald, 2004; Wohr & Soren, 2017), and is thought to integrate emotional and sensory cues (Olson et al., 2007; Pehrs et al., 2017). Therefore, we could speculate that increased activation in response to social alcohol cues in these areas might reflect increased motivation towards socially meaningful stimuli. Still, the evidence for the involvement of brain areas outside of the reward system in substance use needs further exploration. Regarding the absence of specific responses to social alcohol stimuli in reward-related regions, one potential explanation is that alcohol and social cues elicited overlapping activations in these regions (such as the ventral striatum, ACC, and vmPFC, see Figure 1B). This may have resulted in a ceiling effect, leaving little room for an additive effect of social context above and beyond alcohol content.

At the behavioral level, we again expected that the approach bias towards alcohol cues would be stronger in a social context compared with a non-social context. Yet, the current results revealed a general approach bias towards alcohol cues, that was of similar magnitude for social and non-social cues. This suggests that while alcohol cues elicit a stronger approach bias than soda cues, there might be no additive effect of social context in strengthening that bias. The fact that the task instructions were tailored to the alcohol content, i.e. the participants had to either approach or avoid alcohol, might have amplified the focus on the drink and thus mitigated any additional effect of context. In our earlier study using similar social and non-social alcohol stimuli (Groefsema et al., 2016), we observed a comparable main effect of alcohol in the absence of a clear interaction with social context. Our study adds to the literature by replicating our previous findings of a general approach bias towards alcohol (Groefsema et al., 2016) in a young sample of drinkers with varying levels of alcohol use.

Additionally and against our expectations, we found no support for an association between our brain cue-reactivity measures and our behavioral approach measures. This could reflect the fact that the social alcohol cue-reactivity and the stimulus response compatibility tasks engage partly different (brain) mechanisms. More specifically, whereas cue-reactivity (e.g based on the incentive sensitization model of addiction (Robinson & Berridge, 1993, 2001, 2008)), activates the salience and reward brain networks during the anticipation of a reward (Zilverstand, Huang, Alia-Klein, & Goldstein, 2018), approach tendencies (e.g. seen as an automatic response in the dual process models of addiction (Stacy & Wiers, 2010; R. W. Wiers et al., 2007)) require a response and therefore probably also activate the executive network (Zilverstand et al., 2018). This difference between the two measures might explain why they are not correlated in the present data, and further suggest that cue-reactivity and approach biases may be independent mechanisms associated with drinking behaviour.

The present observation that drinking in a social setting was not related to either brain cue-reactivity or behavioural approach biases is in line with previous studies which have failed to pinpoint the origin of individual differences in imitation of drinking. More specifically, these individual differences seem to be unaccounted for by induced stress levels (Larsen, Engels, et al., 2013), engagement between drinking partners (Larsen, Lichtwarck-Aschoff, et al., 2013), or implicit alcohol-related cognitions (Larsen, Engels, et al., 2012). One potential explanation is that environmental factors, such as having peers around, may be stronger predictors of social drinking than individual factors such as brain or behavioral responses to alcohol cues. Supporting this idea, a study by van Schoor et al. (2008) showed that while several personality traits were associated with self-reported daily alcohol consumption or self-reported alcohol-related problems, these personality traits no longer predicted drinking behavior when peers were around. So the company of peers and ‘unwritten’ social norms (Jackson et al., 2014; Teunissen et al., 2012) may have a bigger impact on drinking behaviour than individual traits like the response to alcohol cues.

We believe that the results of this study highlight two further issues that surface if one does not specifically focusses on the effect of the *social* alcohol cues but more on responses to alcohol cues in general. First, it can be questioned what predictive validity approach biases hold, since the approach bias was not related to a measure of real-world drinking. Approach biases are thought to play an important role in (the transition to) heavy drinking (Lindgren et al., 2018; Robinson & Berridge, 2001). Moreover, after training dependent individuals to avoid alcohol cues instead of approaching them, relapse rates decreased (Eberl et al., 2013; Kakoschke, Kemps, & Tiggemann, 2017; Manning et al., 2016; Rinck, Wiers, Becker, & Lindenmeyer, 2018; C. E. Wiers, Ludwig, et al., 2015; C. E. Wiers, Stelzel, et al., 2015; R. W. Wiers, Eberl, Rinck, Becker, & Lindenmeyer, 2011). Yet, such a re-training has not been successful in undergraduate students in terms of reducing approach biases and drinking behaviour (Lindgren et al., 2015). It may be that re-training an approach bias is only be effective in heavy or dependent drinkers and/or among individuals with the motivation to change their behaviour. Despite that we did not re-train approach biases or measure the motivation to change, it could be hypothesized that our sample had indeed a low motivation to change their behavior, as they were mostly students that generally do not see their alcohol consumption as problematic (Vik, Culbertson, & Sellers, 2000) and were not seeking any help. Future studies in young adult drinkers should test the possible moderating role of motivation to change on the link between approach biases and drinking in the real-word.

Second, our findings emphasize the difficulty – and the importance – of relating laboratory measures with drinking in the real-world. Recent studies among adolescent drinkers have shown no relationship between alcohol approach biases and the levels of alcohol consumption measured at different time points with ecological momentary assessments (Janssen, Larsen, Vollebergh, & Wiers, 2015), or have found such relationships only in individuals with weak inhibition skills (Peeters et al., 2013). Moreover, we have previously shown that cognitive biases towards social alcohol cues are not directly related to drinking in the real-world, but only moderate the association between the number of friends present and alcohol use (Groefsema et al., 2016). With regards to cue-reactivity, this was the first study to directly relate brain cue-reactivity with an ecologically valid measure of social drinking. Previous studies that correlated brain activation with a measure of alcohol use revealed a positive association with VS activation, but this was almost always with a self-report measure of alcohol such as problematic drinking or drinking desire (see meta-analysis Schacht et al., (2013)). Collectively, these studies illustrate the difficulty to identify reliable predictors of real drinking behaviour. Alcohol use is a very complex behaviour and can be affected by many different motivations, with each of them possibly explaining a small amount of variance. New analyses techniques such as machine learning can offer promising opportunities allowing researchers to test multiple different determinants of drinking at once. Indeed, it has previously been found that life experiences, neurobiological differences, and personality appear to be the most important factors influencing binge drinking among adolescents (Whelan et al., 2014). Future research should expand this field of research to other samples and types of drinking behaviour.

One of the major strengths of this study is the large sample size which enhances the interpretability of the null findings, as further suggested by our Bayesian statical analyses which generally revealed support for the null hypotheses. Another strength is the ecological validity of our Bar-Lab procedure to measure drinking in a social setting, along with a triangulation approach between brain responses, behavioral responses and actual drinking behavior in a social setting. The integration of such findings within a large sample is still rather unique. A limitation of this study was that only males were included, and most of them were college students, making it difficult to generalize these findings to young adult drinkers in general. Moreover, our sample showed large variation in heaviness of drinking thereby possibly masking effects. Yet, individuals in this age range are specifically known for drinking heavily in social settings and we, therefore, believe it is important to examine such a sample to reveal the underlying mechanisms of social drinking.

In conclusion, our findings show that social alcohol cues elicit specific responses in brain regions (STS and IPL) that have been associated with emotion processing and social cognition rather than reward processing *per se*. Given that these findings are not aligned with our predictions, replication is needed and we prefer to refrain from making strong claims about how these areas might contribute to social alcohol cue-reactivity at the moment. In addition, we found that our young adult heterogenous drinking sample has approach tendencies towards alcohol cues and no correlations between brain cue-reactivity and behavioural approach biases. Furthermore, we found no evidence supporting a relationship between cue-reactivity and an approach bias towards (social) alcohol cues, nor between either of these measures and drinking behavior in a social setting. Despite the fact that laboratory measures of cue-reactivity and approach biases were not related to drinking in a social setting in the current study, we would like to encourage future studies to continue to include a measure of real-world drinking in combination with, for example, machine learning analyses, in order to strengthen the predictive validity of research in the laboratory to drinking behaviour in the real-world.

## ACKNOWLEDGEMENTS

This research did not receive any specific grant from funding agencies in the public, commercial, or not-for-profit sectors.

## AUTHOR CONTRIBUTIONS

All authors contributed to the design of the study. MG collected the data. MG, GS, GM and ML performed data analyses. MG, GM, GS and ML wrote the first draft of the manuscript. RE and JC edited the manuscript.

## DATA ACCESIBILITY STATEMENT

All unthresholded T-maps resulting from the fMRI analyses can be accessed at https://neurovault.org/collections/IVCNOFBQ/. Data are available from the corresponding author on request.

## ABBREVIATIONS

ACC: anterior cingulate cortex
AUDIT: Alcohol Use Disorders Identification Test
BF: Bayes Factor
BOLD: Blood oxygen level-dependent
DSM-IV: Diagnostic and Statistical Manual of Mental Disorders – 5
fMRI: functional Magnetic Resonance Imaging
FWE: family-wise error
ICA: independent component analysis
AROMA: automatic removal of motion artifacts
IPL: inferior parietal lobe
MRI: Magnetic Resonance Imaging
NA: Non-Social Alcohol
NS: Non-Social Soda
PCA: Principal Component Analysis
SA: Social Alcohol
SACE: Social-Alcohol Cue-Exposure
SPSS: Statistical Package for Social Sciences
SRC: Stimulus-Response Compatibility
SS: Social Soda
STS: superior temporal sulcus
vmPFC: ventromedial prefrontal cortex
VS: ventral striatum

