## Supplementary Material for "Cue-reactivity and approach bias to social alcohol cues and their association with drinking in a social setting in young adults"

Figure S1: Flow chart of the entire data collection

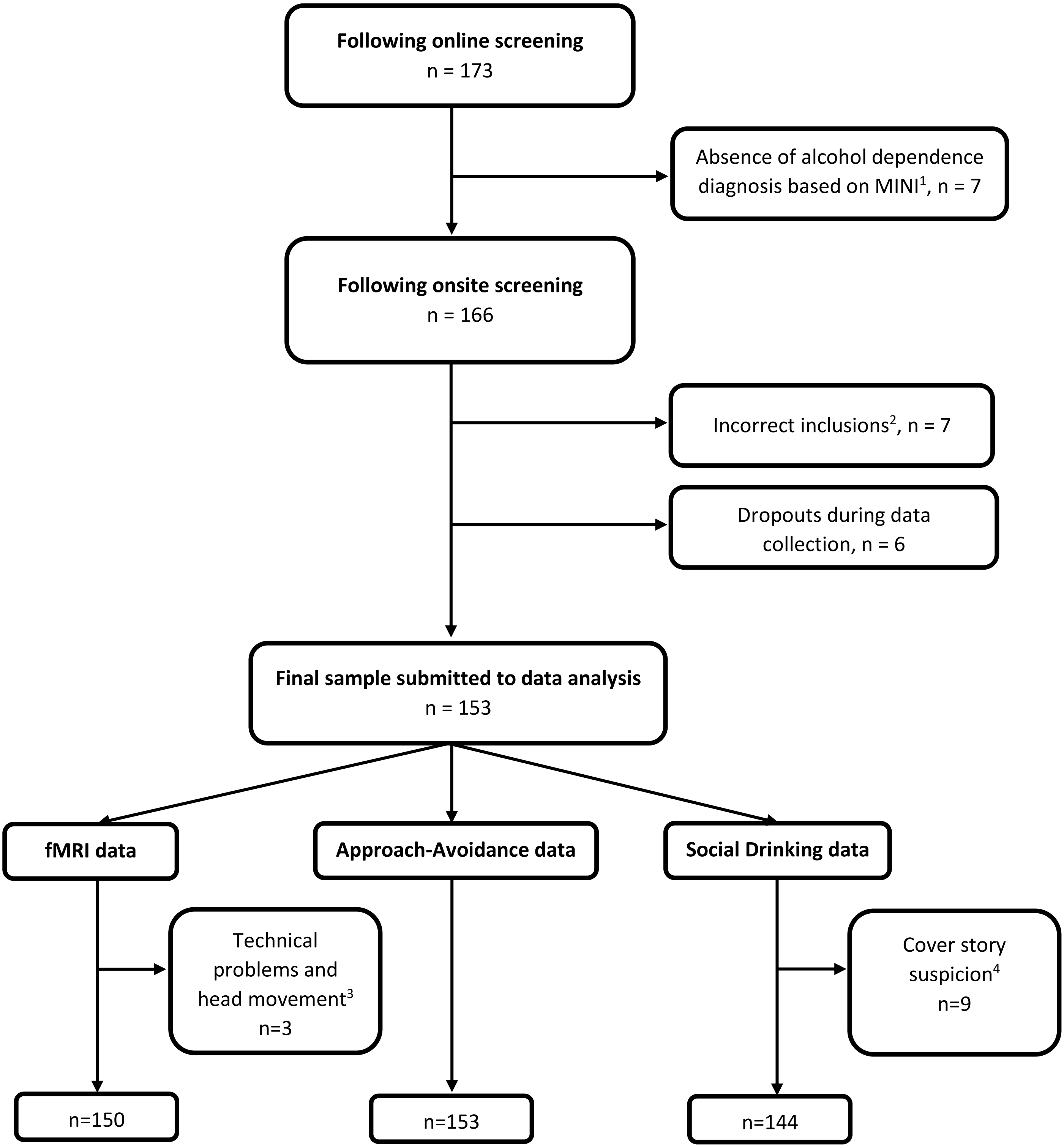

*[1]In the context of the larger project, a severe group of "dependent drinkers" was defined based on AUDIT scores, weekly drinking scores and DSM criteria for alcohol dependence using a MINI interview. Some participants who showed signs of problematic and heavy alcohol use during online screening did not meet the criteria for alcohol dependence following the onsite screening. We decided to exclude these participants, as we deemed that this inconsistency between self-report and DSM-based measures made their categorization uncertain.*

[2] *Incorrect inclusions did not meet the combined requirement for the AUDIT and weekly drink score to be included in the "light drinkers" or " at-risk drinkers" group, also in the context of the larger project. The data from these 7 participants were discarded before performing any data analysis.*

[3] *For one participant, data from the fMRI session was missing, because we were unable to correct for his impaired vision. For another two participants, the fMRI data were discarded because they did not pay attention during the SACE-task as they did not respond to the control pictures using a button-press.*

[4] *Nine individuals appeared to be aware of the true purpose of the Bar-Lab sessions, and were therefore excluded from the calculations on the social drinking data.*

Table S1: Overview of all collected data in the study

| WHEN | WHAT | HOW |
|---|---|---|
| **SCREENING** | | |
| | Alcohol use disorder | AUDIT |
| | Drinks last week | TLFB |
| | Gender | Male/female question |
| | Age | Open question |
| **MINI INTERVIEW (for the dependent group only)** | | |
| | Alcohol dependence diagnosis | MINI |
| | Other substance use information | MINI |
| **BEHAVIOURAL SESSION 1** | | |
| | Education | Multiple choice |
| | Age of first alcohol consumption | Open question |
| | Frequency of drinks in the last 4 weeks | Multiple choice |
| | Binge drinking episodes in last 4 weeks | Multiple choice |
| | Location of alcohol consumption | Multiple choice |
| | Drinking motives | DMQ-R |
| | Ever drank alcohol | Yes/No question |
| | Number of hours since last drink | Open question |
| | Ever smoked in life | Yes/No question |
| | Current smoker | Yes/No question |
| | Number of hours since last cigarette | Open question |
| | Smoking severity | FTND |
| | Ever used drugs (sleeping pills/cannabis/cocaine/ecstacy/amphetamine/hallucinogens/opiates), if yes how many times | Open question |
| | Impulsivity | BIS11 |
| | State-trait Anxiety | STAI |
| | Depression | BDI |
| | Drinking Urge | DAQ |
| | Imitation of alcohol use | Number of drinks consumed in Bar-lab |
| | Confederate liking ratings | 1-9 scales |
| | Anxiety before/during/after session | Multiple choice |
| | Suspicion checks about study goals | Open questions |
| **BEHAVIOURAL SESSION 2** | | |
| | Number of hours since last drink | Open question |
| | Number of hours since last cigarette | Open question |
| | Delay Discounting | Delay discounting task |
| | Drinking Urge | DAQ |
| | Imitation of alcohol use | Number of drinks consumed in Bar-lab |
| | Confederate liking ratings | 1-9 scales |
| | Anxiety before/during/after session | Multiple choice |
| | Suspicion checks about study goals | Open questions |

*Table S1: Overview of all collected data in the study (continued)*

| WHEN | WHAT | HOW |
|---|---|---|
| **FMRI SESSION** | | |
| **WHEN** | **WHAT** | **HOW** |
| | Number of hours since last drink | Open question |
| | Number of hours since last cigarette | Open question |
| | Height/Weight | Open question |
| | Drinking Urge | DAQ |
| | Drinking Self-Efficacy | DRSEQ |
| | Brain responses to social alcohol cues | Social Alcohol cue Reactivity task |
| | Brain responses to anticipating and receiving beer | Beer Incentive Delay task (current paper) |
| | Approach/Avoidance of (social) alcohol pictures | Stimulus-Response Compatibility task |
| **FOLLOW UP BASELINE** | | |
| | Alcohol use disorder | AUDIT |
| | Average number of drinks for each day of the week | Weekly drinking |
| | Drinking motives | DMQ-R |
| **FOLLOW UP WITH ECOLOGICAL MOMENTARY ASSESSMENT (14 DAYS)** | | |
| | Number of (non)alcohol units consumed day before | Open question |
| | Location of (non)alcohol consumption day before | Multiple choice |
| | People with whom (non)alcohol was consumed with day before | Multiple choice |

*Table S2: Whole brain activations for the different contrasts in the SACE task*

| Brain area | Hemisphere | Cluster size | MNI coordinates peak voxel | | | T-value | p-value |
|---|---|---|---|---|---|---|---|
| | | | x | y | Z | | |
| Interaction ((SA>SS)-(NA>NS)) | | | | | | | |
| **Superior Temporal Sulcus** | **R** | **104** | **54** | **-4** | **-11** | **4.75** | **0.005** |
| Superior Temporal Sulcus | R | | 51 | 5 | -14 | 4.17 | |
| Superior Temporal Sulcus | R | | 48 | -10 | -8 | 4.02 | |
| **Superior Temporal Sulcus** | **L** | **103** | **-51** | **8** | **-8** | **4.48** | **0.006** |
| Superior Temporal Sulcus | L | | -60 | 5 | 19 | 3.80 | |
| Superior Temporal Sulcus | L | | -54 | -1 | -20 | 3.45 | |
| **Inferior Parietal Lobe** | **L** | **66** | **-60** | **-28** | **25** | **4.16** | **0.036** |
| Alcohol > Soda ((SA+NA)-(SS+NS)) | | | | | | | |
| **Visual Cortex** | **L** | **2359** | **-15** | **-88** | **-11** | **16.37** | **0.000** |
| Visual Cortex | R | | 18 | -94 | 1 | 13.89 | |
| Visual Cortex | L | | -18 | -100 | 7 | 13.60 | |
| **Superior Frontal Gyrus** | **L** | **430** | **-3** | **35** | **-5** | **5.70** | **0.000** |
| Anterior Cingulate Cortex | L | | -3 | 17 | -6 | 4.84 | |
| Anterior Cingulate Cortex | L | | -12 | 41 | -11 | 4.43 | |
| **ventral medial Prefrontal Cortex/ Subgenual Frontal Cortex** | **L/R** | **424** | **0** | **11** | **55** | **5.66** | **0.000** |
| ventral medial Prefrontal Cortex/ Subgenual Frontal Cortex | L/R | | 0 | 23 | 31 | 5.55 | |
| Superior Frontal Gyrus | L | | -3 | 17 | 40 | 5.42 | |

*Table S2: Whole brain activations for the different contrasts in the SACE task*

| | | | | | | | |
|---|---|---|---|---|---|---|---|
| **Superior parietal gyrus** | **L** | **307** | **-15** | **-52** | **10** | **4.89** | **0.000** |

| | | | | | | | |
|---|---|---|---|---|---|---|---|
| Superior parietal gyrus | L | | -6 | -61 | 13 | 4.73 | |
| Superior parietal gyrus | R | | 18 | -52 | 13 | 4.58 | |
| Social > Non-Social ((SA+SS)-(NA+NS)) | | | | | | | |
| **Cuneus** | **R** | **14291** | **12** | **-88** | **1** | **26.22** | **0.000** |
| Lingual Gyrus | L | | -9 | -88 | -5 | 24.72 | |
| Lingual Gyrus | R | | 12 | -85 | -8 | 24.30 | |
| **Middle frontal gyrus** | **R** | **908** | **45** | **14** | **31** | **11.93** | **0.000** |
| Inferior Frontal Gyrus | R | | 57 | 29 | 19 | 11.31 | |
| Inferior frontal gyrus | R | | 48 | 23 | 22 | 10.78 | |
| **Ventral medial Prefontal Cortex/Posterior orbital gyrus** | **R** | **69** | **33** | **35** | **-14** | **8.55** | **0.047** |
| ventral medial Prefrontal Cortex /Posterior orbital gyrus | R | | 30 | 29 | -2 | 4.52 | |
| **Superior Frontal Gyrus** | **L** | **109** | **-6** | **14** | **49** | **6.19** | **0.008** |
| Superior Frontal Gyrus | R | | 6 | 14 | 49 | 5.97 | |

*Note:* Reported coordinates correspond to the three highest peak voxels more than 8 mm apart in each cluster. P-values are cluster level FWE-corrected.
